# Dynamic B_0_ field shimming for improving pseudo-continuous arterial spin labeling at 7 Tesla

**DOI:** 10.1101/2024.07.22.604544

**Authors:** Yang Ji, Joseph G. Woods, Hongwei Li, Thomas W. Okell

**Author notes:** Correspondence: Joseph G. Woods, Ph.D., Wellcome Centre for Integrative Neuroimaging, FMRIB, Nuffield Department of Clinical Neurosciences, University of Oxford, Oxford, UK.

## Abstract

**Purpose:** B_0_ inhomogeneity within the brain-feeding arteries is a major issue for pseudo-continuous arterial spin labeling (PCASL) at 7T because it reduces labeling efficiency and leads to a loss of perfusion signal. This study aimed to develop a vessel-specific dynamic B_0_ field shimming method for 7T PCASL to enhance labeling efficiency by correcting off-resonance in the arteries within the labeling region.

**Methods:** We implemented a PCASL sequence with dynamic B₀ shimming at 7T that compensates for B_0_ field offsets at the brain-feeding arteries by updating linear shimming terms and adding a phase increment to the PCASL RF pulses. Rapidly acquired vessel-specific B₀ field maps were used to calculate dynamic shimming parameters. We evaluated both 2D and 3D variants of our method, comparing their performance against established global frequency offset and optimal-encoding-scheme (OES)-based corrections. Cerebral blood flow (CBF) maps were quantified before and after corrections. CBF values from different methods in the whole brain, white matter, and grey matter regions were compared.

**Results:** All off-resonance correction methods significantly enhanced perfusion signals across the brain. The proposed vessel-specific dynamic B₀ shimming method improved labeling efficiency while maintaining optimal static shimming in the imaging region. Perfusion-weighted images demonstrated the superiority of 3D dynamic B₀ shimming method compared to global or 2D-based correction approaches. CBF analysis revealed that 3D dynamic B₀ shimming significantly increased CBF values relative to the other methods.

**Conclusion:** Our proposed dynamic B_0_ shimming method offers a significant advancement in PCASL robustness and effectiveness, enabling full utilization of 7T ASL’s high sensitivity and spatial resolution.

## Introduction

Arterial spin labeling (ASL) offers a non-invasive method for quantifying tissue perfusion by leveraging endogenous blood water protons as a contrast agent (1,2). This technique has gained widespread adoption in both research and clinical settings due to its safety and ability to provide detailed information about blood flow to various organs and tissues (3–6). In addition to conventional ASL techniques that rely on a long post-labeling delay (PLD) to visualize tissue perfusion, ASL can also be used for acquiring images with short PLDs (7,8). These short-delay ASL images capture the labeled blood primarily within the arteries, providing angiographic contrast. Pseudo-continuous arterial spin labeling (PCASL) stands out among various ASL methods due to its superior labeling efficiency to continuous ASL (CASL), higher signal-to-noise ratio (SNR) than pulsed ASL (PASL), and greater compatibility with modern clinical scanners, making it ideal for clinical MRI applications (9,10). Unlike CASL, which requires continuous radiofrequency (RF) pulses that can be technically challenging due to power and duty cycle limitations, PCASL employs a series of closely spaced low flip-angle RF pulses and gradients, effectively removing these challenges. ASL typically involves two acquisitions: a labeling scan, where blood water magnetization is inverted, and a control scan, where magnetization remains unaltered. Perfusion information is then extracted by subtracting the labeled scan from the control scan. While this method effectively captures tissue perfusion, the low blood water concentration and T_1_-decay of the labeled blood inherently results in a low signal-to-noise ratio (SNR). This characteristic presents significant challenges in achieving high-quality perfusion measurements. Traditionally, improving SNR in MRI has relied on increasing signal averages. However, this approach extends scan duration, potentially leading to subject discomfort and increased motion artifacts, which can degrade image quality. Ultra-high field (UHF) MRI has emerged as a promising avenue to boost ASL’s SNR due to several factors(11). Firstly, the intrinsic SNR exhibits a theoretical dependence on the static magnetic field strength (B_0_) following a power law relationship, scaling with B₀^1.94 ± 0.16^(12). Secondly, the extended ASL tracer lifetime (i.e., blood T_1_) at a higher field indirectly contributes to increased SNR in ASL(13,14). However, implementing ASL sequences at UHF presents challenges, primarily due to specific absorption rate (SAR) limitations as well as B_0_ field and transmit B_1_ field (B_1_^+^) inhomogeneities, especially in the neck. These inhomogeneities can substantially reduce labeling efficiency, thereby reducing the perfusion signal. In severe cases where there are significant offsets from the nominal B₀ in PCASL, the perfusion signal can shrink to zero, or the labeling and control scans may even be completely reversed. It’s crucial to recognize that B₀ field inhomogeneities vary considerably across different vessels, individuals, and even repeated scans, influenced by factors such as static shimming, subject positioning, the placement of the labeling region, and individual anatomical differences(15,16). This variability poses a substantial challenge in achieving consistent cerebral blood flow (CBF) measurements with ASL at UHF, thereby impacting its accuracy and reproducibility.

(17,18) Various compensation methods have been explored to address this challenge. A straightforward approach using static shimming applied to both labeling and imaging regions partially alleviates the issue (19), but residual inhomogeneity remains problematic. Additionally, spatial separation between the labeling and imaging regions presents an additional challenge for shimming and can lead to suboptimal shimming settings for both regions, ultimately compromising image quality (i.e., distortion/signal dropout for EPI) and labeling efficiency. Unbalanced PCASL with optimized parameters has been proven to offer improved performance for off-resonance effects, but significant signal loss persists at larger phase offsets, limiting its effectiveness(20). Multi-phase PCASL (MP-PCASL) addresses frequency offset issues by acquiring data at multiple RF phase increments instead of the conventional two (0 and π)(21). This enables a more precise fit of voxel data to predicted response curves, reducing phase mismatches and resulting in more accurate and reliable CBF measurements. However, MP-PCASL suffers from lower SNR or increased scan times, which may limit its widespread adoption in research and clinical settings. A strategy was initially proposed to address B_0_ inhomogeneity in the labeling region of PCASL scans by applying an additional z-gradient and RF phase adjustment (15). This method uses a coronal field inhomogeneity map to identify frequency offsets for arteries at various z-axis locations. Based on these frequency offsets and corresponding z-axis location information, a z-gradient term and an additional RF phase term are calculated to compensate for z-direction B_0_ inhomogeneity and residual global frequency offset, respectively. While this approach effectively mitigates B_0_ inhomogeneity, it does not account for in-plane variations. This limitation becomes crucial when arteries within the same slice have significantly different off-resonance values. OptPCASL offers a more comprehensive solution by addressing B_0_ field inhomogeneity, including both the global frequency offset and the local variations between the labeled feeding arteries (22). This approach accounts for the global phase error, which affects all feeding arteries uniformly, and the local phase errors specific to individual feeding arteries. It achieves this by incorporating an additional compensation RF phase term into both the tag and control conditions of the ASL sequence, and by applying extra in-plane gradient pulses between the RF tagging pulses. A very similar method has been developed at 7T (23), and both methods utilize an MP-PCASL pre-scan to measure the phase tracking errors that are used to estimate the RF phase increment and transverse gradients required, which extends the total scan time. An optimized encoding scheme (OES)-based method has also been proposed to correct for off-resonance at an arbitrary number of vessel locations using a rapidly acquired field map of the labeling plane (16,24). Transverse gradients and RF phase increments are then calculated using the Fourier-based OES method to compensate for the offset at each vessel location. While the three aforementioned methods effectively address frequency offset of B_0_ field variations across the vessels, they do not account for through-plane B_0_ field inhomogeneity. This could be problematic if the frequency variations of arteries along the through-plane are significant.

In this work, we propose a novel dynamic B_0_ shimming method to improve PCASL labeling efficiency at 7T by addressing B_0_ field inhomogeneity directly. Unlike conventional static B_0_ shimming, which maintains constant shimming settings throughout the entire scan, this method dynamically adjusts the linear (gradient) shimming currents between the labeling period and the imaging readout, providing optimal B_0_ homogeneity for both labeling and imaging. This proposed correction method addresses both in-plane and through-plane B_0_ field inhomogeneities within the labeled arteries. By targeting just the B_0_ field inhomogeneity within the feeding arteries in the labeling region, rather than the entire labeling region (including static tissue), this method achieves significantly enhanced correction effectiveness. We compared our proposed method to existing correction techniques for B_0_ field inhomogeneity in PCASL at 7T, demonstrating its effectiveness and superiority. This study highlights the feasibility of our approach, achieving high-quality PCASL perfusion data in the whole human brain at 7T.

## Methods

### PCASL implementation with dynamic B_0_ shimming

The proposed PCASL sequence diagram with dynamic B_0_ shimming is illustrated in Figure 1. The PCASL tagging/control module consists of a train of discrete slice-selective RF pulses applied at intervals of Δτ. To correct B_0_ inhomogeneity within the labeling region, the system dynamically adjusts first-order shimming parameters during the labeling period. Since first-order shimming relies on modifying the linear x, y, and z gradients, these adjustments introduce offsets to the gradient amplitudes (i.e, Δ*G_x_*, Δ*G_y_*, Δ*G_z_*) compared to the initial values throughout the PCASL pulse train. The residual global frequency offset (Δ*f_Glob_*) is corrected by adding additional phase increments to the PCASL RF pulses. Thus, in addition to 0 and π phase, an additional ϕ*_off_* that includes both phase corrections for global off-resonance and the off-isocenter of the labeling plane is added into the RF phases for both control and tagging, which is given by (25):

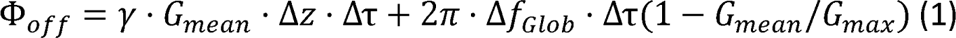

**Figure 1.**
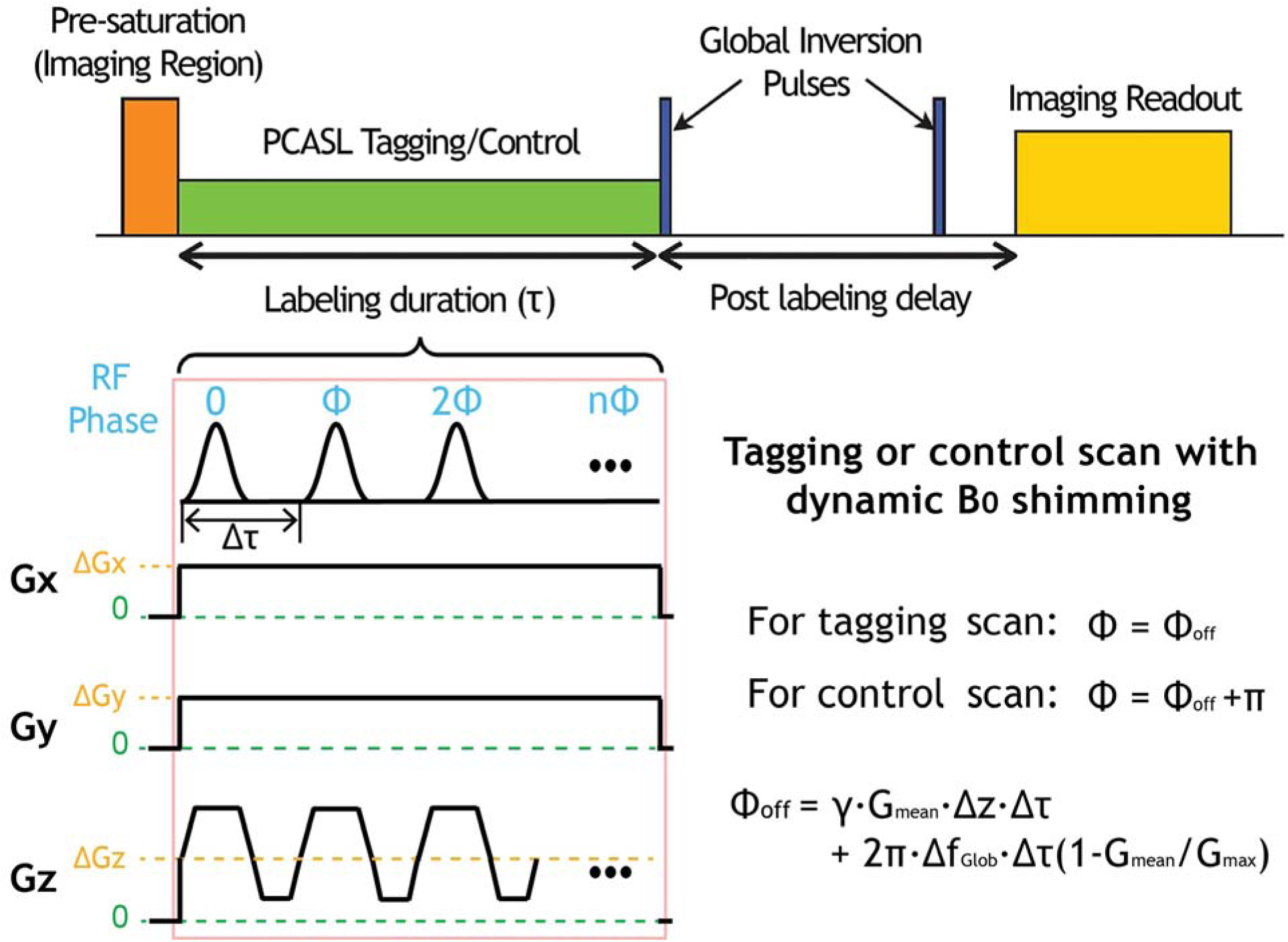
This diagram illustrates the proposed PCASL sequence incorporating dynamic B_0_ shimming for field inhomogeneity correction. The PCASL module consists of a train of discrete slice-selective RF pulses every Δτ. To compensate for B_0_ inhomogeneity, additional gradients with constant amplitude are applied along three axes during the PCASL pulse train. Additionally, a residual global frequency offset (Δ*f_Glob_*) is corrected by incorporating specific phase increments into the PCASL RF pulses. Note that ϕ*_off_* include both phase corrections for global off-resonance and the off-isocenter of the labeling plane.

where G_means_ the mean gradient, G_max_ is the maximum gradient, Δz the offset of the nominal labeling plane, Δf_Glob_ is the residual global frequency offset. The calculation of Δ*f_Glob_*, along with the parametersΔG_x_, ΔG_y_, and ΔG_z_ are detailed in the following subsection. Note that the (1 – G_mean_ / G_max_) correction term accounts for the small shift in the labeling plane location due to the global frequency offset.

### Vessel-specific dynamic B0 shimming

The required offset amplitudes of the gradients (Δ*G_x_*, Δ*G_y_*, Δ*G_z_*) for dynamic B_0_ shimming and the residual global frequency offset (Δ*f_Glob_*) can be estimated for specific vessels using field maps that encompass the labeling region. This estimation is achieved by solving the following equation:

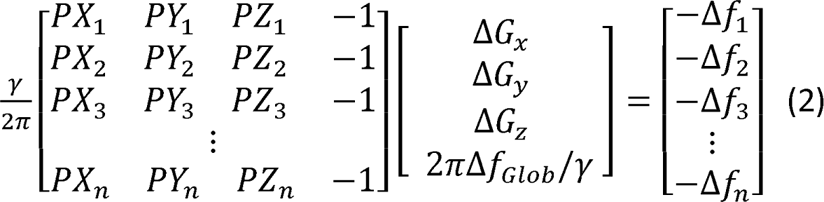

where [PX_i_, PY_i_, PZ_i_] and Δf_i_ represent the location of the i-th vessel voxel within the 3D shimming region and the corresponding global frequency offset, respectively. Since the labeling efficiency is only affected by B_0_ homogeneity of the flowing blood inside the brain-feeding arteries within the labeling region, we restrict the shimming optimization area to small ROIs encompassing only the relevant voxels. This targeted method allows for much more efficient B_0_ shimming by focusing solely on the relevant vessel voxels within the labeling region, excluding static tissues and air that do not impact the labeling process. Since there are only four unknown parameters and typically many voxels within the ROIs, this is a well-conditioned problem that can be solved simply by taking the pseudoinverse of the matrix on the left-hand side of Eq. 2. Additionally, dynamic B_0_ shimming can also be performed in two-dimensions (2D) when ignoring through-plane B_0_ variations:

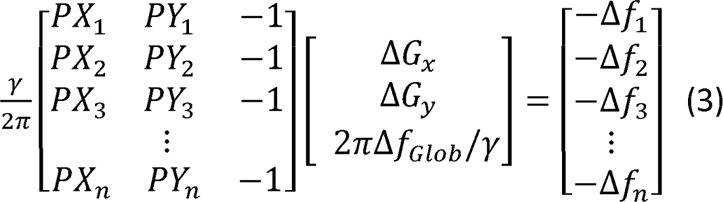

Under such conditions the shimming gradient’s offset along z-direction should be set to zero. Similarly, if the through-plane B_0_ variations are only considered and the offset amplitudes for the transverse plane gradients (x and y) are instead set to zero, this method essentially reduces to the one proposed by Jahanian et al. (15), as mentioned above.

Figure 2 illustrates the workflow of PCASL at 7T with dynamic B_0_ shimming. A three-dimensional (3D) time-of-flight (TOF) scan is performed to select the PCASL labeling plane location, positioned in the inferior-superior direction at the middle of the V3 section of the vertebral arteries as in previous studies(26) to allow whole-brain perfusion measurements. To acquire B_0_ field inhomogeneity information within the labeling region, a subsequent field map sequence is performed with its imaging volume centered on the labeling plane. It is important to note that the static B_0_ shimming volume of the field mapping sequence should match that of the subsequent PCASL scan. Vessel ROIs are manually drawn to allow frequency offsets and voxel locations to be extracted from the field map data and the dynamic B_0_ shimming-related parameters to be calculated. These parameters are then applied during the PCASL scan, which utilizes an identical static B_0_ shimming configuration as the field mapping sequence.

**Figure 2.**
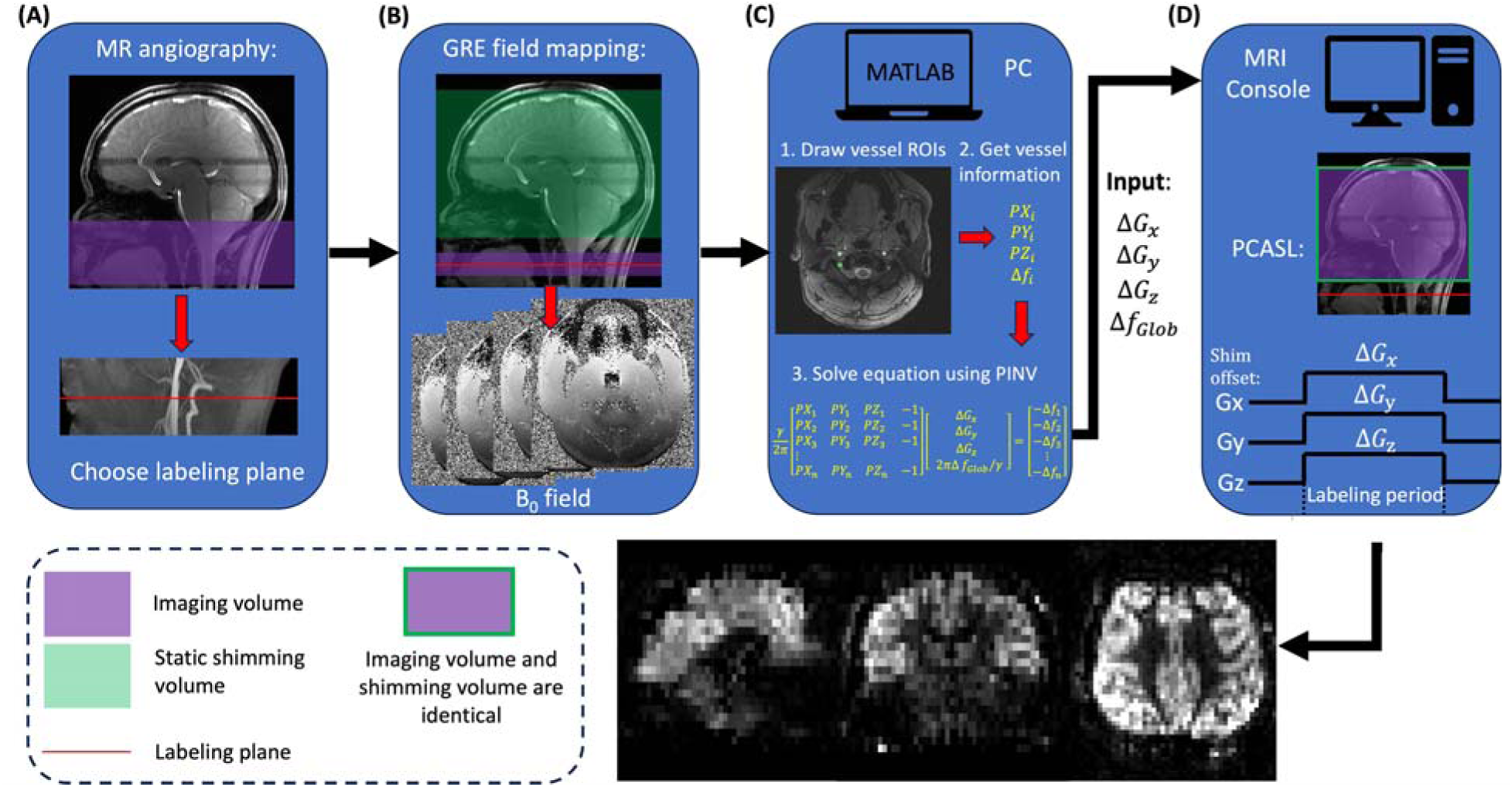
Workflow of using dynamic B_0_ shimming in PCASL **(A)** First step: perform a TOF sequence to select the optimal labeling plane (placed at the middle of the V3 section of the vertebral arteries). **(B)** Second step: conduct a 2D multi-slice GRE field mapping sequence centered at the labeling plane, covering the labeling region, to map the frequency offsets of the vessel voxels within this region. Note that the fieldmap field-of-view is assumed to cover the region over which blood labeling occurs. **(C)** Third step: manually draw the vessel ROIs within the labeling region in MATLAB, then pass the frequency offset and corresponding position in the scanner coordinate system to construct the equation. Finally, use MATLAB’s PINV function to calculate the required gradient offsets and global frequency offset. **(D)** Fourth step: input the calculated values into the MRI console and run the PCASL with the static B_0_ shimming volume matching that of the GRE field mapping sequence, ultimately generating high-quality perfusion images.

### In vivo data acquisition

Data were acquired on a MAGNETOM 7T Plus scanner (Siemens Healthineers, Erlangen, Germany) equipped with an 8Tx/32Rx head coil. Six healthy volunteers (aged 23 to 34) were scanned under a technical development protocol with approval from local ethical and institutional review boards. A 3D multi-slab TOF angiography sequence with voxel size of 0.8 × 0.8 × 1.3 mm³ was conducted in the neck, enabling the selection of an optimal labeling plane. A rapid 2D multi-slice dual-echo GRE field mapping sequence was performed for collecting the frequency offset information of the vessel voxels within the labeling region. The main sequence parameters were as follows: TE1 = 4.08 ms, TE2 = 5.1 ms; number of slices = 5; FOV = 220 × 220 mm^2^, TR = 40 ms; in plane spatial resolution = 0.7 × 0.7 mm^2^; slice thickness = 2 mm, scan time = 26 s. To calibrate the transmit voltage for the brain, a 3DREAM B_1_^+^-mapping sequence (27) was performed with the imaging volume including the whole brain and the neck. The average voltage from the central slice of the brain was then updated in the system. To mitigate the B_1_^+^ drop-off at the labeling plane, the nominal flip angle of the B_1_^+^ pulse for labeling was adjusted by multiplying the target flip angle by a factor of V_brain_/V_vessel_, where V_brain_ represents the average voltage across the central slice of the brain, and V_vessel_ represents the average voltage across the labeled vessel voxels.

To evaluate the performance of the proposed dynamic B_0_ shimming method for correcting B_0_ inhomogeneity in PCASL at 7T, whole-brain perfusion images using PCASL with existing correction methods (global frequency offset correction(28), OES-based(16)) and without correction were acquired at 7T. A PCASL sequence with a 2D gradient-echo EPI readout was employed, utilizing low-SAR optimization(29). The labeling parameters were as follows: labeling RF pulse = Hann-shaped pulse, RF duration = 530 μs, RF separation = 1060 μs, labeling duration = 1400 ms, target RF flip angle = 9.8° (with the nominal flip angle value on the console adjusted to compensate for the B_1_^+^ drop-off in the neck, as described above), mean gradient (G_mean_) = 0.20 mT/m, maximum gradient amplitude (G_max_) = 5.5 mT/m, PLD = 1800 ms. The EPI readout employed the following parameters: FOV = 220 × 220 mm², voxel size = 3.4 × 3.4 × 5 mm³, matrix size = 64 × 64, FA = 90°, phase partial Fourier factor = 6/8, no in-plane acceleration, TE/TR = 13.0/5200 ms, number of slices = 24, and bandwidth = 2004 Hz/pixel. For the conventional tag/control mode, a total of 41 images, including the initial calibration (M_0_) image and alternating control and tagging images, were acquired, with a total acquisition time of 3:35 min. In addition, PCASL images of 24 axial slices with high spatial resolution were obtained, with voxel size of 2.0 × 2.0 × 4 mm³, TE/TR = 13.0/6300 ms, an in-plane GRAPPA(30) acceleration factor of 2, and a scan time of 4:32 min. To enable distortion correction of the EPI images before analysis, field maps covering the perfusion imaging volume were acquired with a static shimming setting matching that of the EPI readout (voxel size: 2.0 × 2.0 × 2.0 mm³, TR: 620 ms, ΔTE: 1.02 ms, acquisition time: 2:02 min). Additionally, T1-weighted structural images using MPRAGE were acquired with a voxel size of 0.7 × 0.7 × 0.8 mm³ to serve as an anatomical reference.

### In vivo data processing

In vivo data were analyzed with the combined use of MATLAB (MathWorks, Natick, MA) and FSL (Oxford, UK) software(31). The vessel ROIs were manually drawn on the magnitude image from the field mapping sequence. The frequency offset of the vessel voxels was calculated by 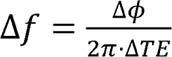, where Δ*TE* is the TE difference, and Δϕ is the phase difference between two echoes. Δϕ was computed by taking the angle of a magnitude-weighted complex sum across channels: 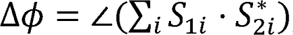 where *S*1*_i_* and *S*_2*i*_ the complex signals from channel *i* for the first and second echo, respectively, and ∗ denotes the complex conjugate operator. Given a Δ*TE* of 1.02 ms, which resulted in an approximate dynamic range of ±490 Hz, and considering that the frequency offset of the vessel voxels was within the range of approximately ±200 Hz, phase unwrapping was not applied during the calculation of the field map for the vessel voxels.

Pre-processing of T1-weighted images from MPRAGE was performed using FSL’s fsl_anat tool, which included bias correction, brain extraction, and tissue-type segmentation. Whole brain B_0_ field maps for distortion correction were generated using FSL’s fsl_prepare_fieldmap tool. All PCASL raw images were motion corrected to the first (calibration) volume image as a reference using FSL’s MCFLIRT(32) tool. Perfusion-weighted images (PWI) were generated by computing the difference between the tag and corresponding control images, and then averaging this difference across all repetitions. The software OXASL (33,34) (https://github.com/physimals/oxasl), which performs Bayesian analysis of arterial spin labeling MRI data, was used for further CBF quantification. Field maps, pre-processed T1-weighted images, and motion-corrected PCASL images were provided as input to OXASL. ROIs for white matter (WM) and grey matter (GM) in structural space were generated using FSL_anat. OXASL subsequently registered these ROIs into ASL space, resulting in ROIs in ASL space.

### Bloch simulation and statistical analysis

We conducted a Bloch simulation to determine the z-direction extent of the labeling region in PCASL using low SAR parameters for this study. The simulation was performed using MATLAB code, which is available on https://github.com/tomokell/bloch_sim(35) using the same parameters as those for in vivo acquisitions. To simplify interpretation of the labeling region, we assumed T_1_ = T_2_ = ∞ for this simulation. Regional CBF values were calculated within the defined ROIs. Paired t-tests were employed to compare quantitative CBF values obtained from PCASL scans with various correction methods. A value of P < 0.05 was considered statistically significant. All data are reported as mean ± standard deviation (SD).

## Results

To compare the influence of two different static B_0_ shimming settings on field homogeneity during the labeling and imaging periods of PCASL, a whole brain PCASL scan and a field mapping scan at the labeling plane were performed on the healthy subjects. Figure 3A illustrates the two different static B_0_ shimming setups, one with the shimming optimized only for the imaging volume, while the other was optimized for the entire imaging volume with an extended bottom edge to include the labeling plane. Field maps of the labeling plane (Figure 3B) showed a slightly larger frequency offset when shimming was restricted to the imaging volume compared to combined imaging and labeling plane shimming, indicating static B_0_ shimming that covers both regions is better for the labeling process of PCASL, although considerable off-resonance remains. Figure 3C shows a representative slice acquired using PCASL-EPI with the two different static shimming settings, superimposed with the outline of a structural image obtained with MPRAGE. The EPI images from the shimming setting that covers only the imaging region show less distortion, indicating that the typical shimming approach for PCASL at 7T that covers both the imaging region and labeling plane is suboptimal for the imaging readout and background suppression pulses. Consequently, the shimming setup that covers only the imaging volume was used in subsequent PCASL experiments with the dynamic B0 shimming correction method.

**Figure 3.**
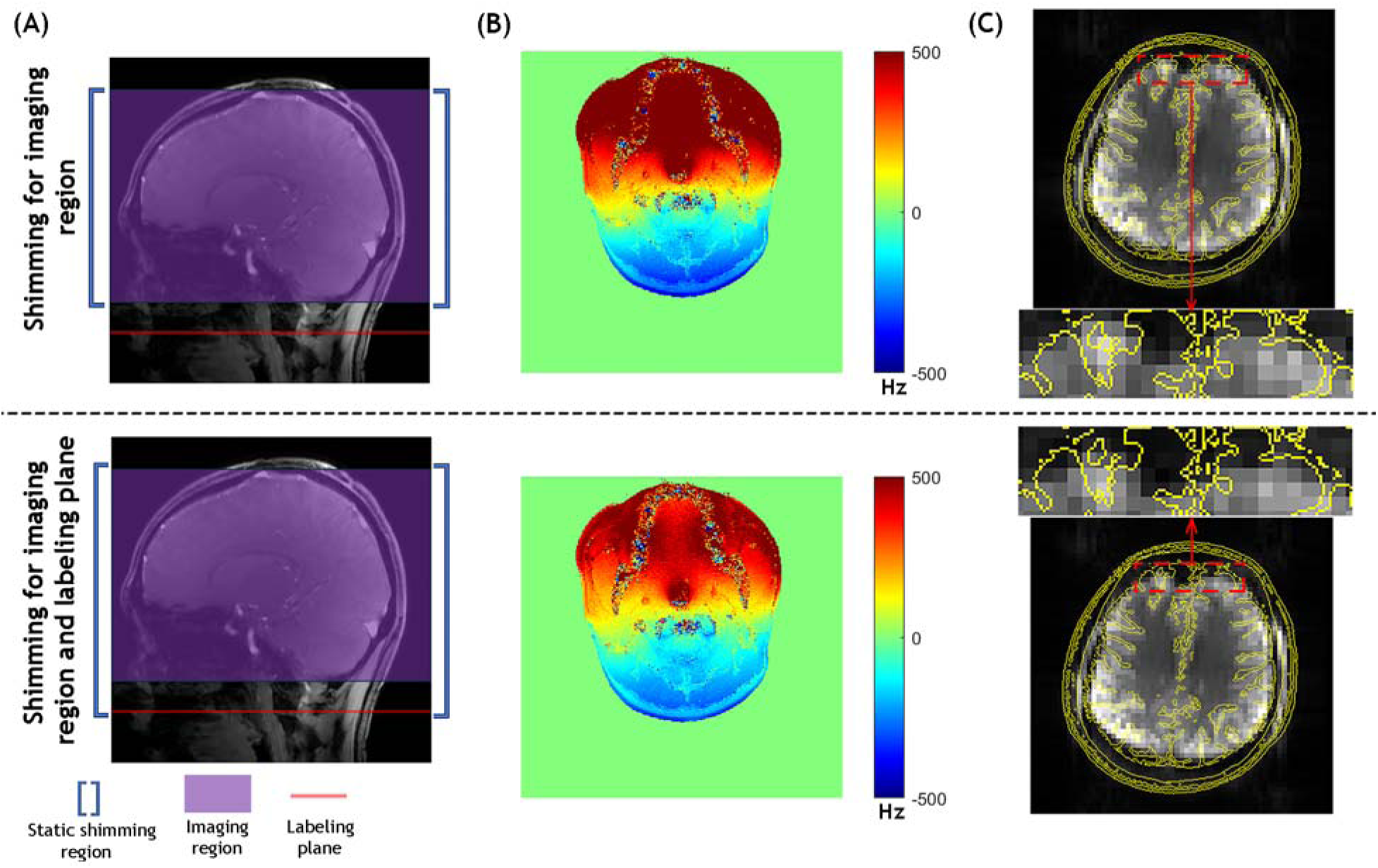
Comparison of two static B_0_ shimming approaches for PCASL **(A)** Static shimming regions: imaging-specific (top) vs. combined imaging and labeling plane (bottom). **(B)** Representative field maps at the labeling plane for both shimming settings, showing there is still considerable off-resonance present at the vessel locations within the labeling plane even when it is included in the static shimming optimization region. **(C)** MPRAGE structural image outline (yellow) overlaid on PCASL-EPI whole brain image slice (grey) for both shimming approaches. This overlay enables the visualization of EPI image distortion, demonstrating improved B_0_ field homogeneity when static B_0_ shimming only for the imaging region.

PCASL literature often uses “labeling plane” interchangeably with “labeling region,” even though the labeling region encompasses a larger volume around the plane where the blood is labeled. To avoid confusion in this study, we strictly define the labeling region as the area where blood undergoes longitudinal magnetization inversion, transitioning from a positive threshold value to a negative threshold value, and the labeling plane as the center of the defined labeling region. It is important to note that while the labeling plane has a precise position, the boundaries of the labeling region are not explicitly defined in existing literature. Additionally, the thickness of the labeling region is influenced by the specific labeling parameters used. Figure 4A illustrates the spatial relationship between the labeling region and the labeling plane of PCASL, along with the four main brain-feeding arteries labeled within the labeling plane (Figure 4B). Figure 4C depicts the simulated evolution of the longitudinal magnetization (M_z_) value along the z-axis during the PCASL labeling process using our low SAR parameters. In our study, ±70% thresholds of equilibrium magnetization were used to define the labeling region’s boundaries, which resulted in a labeling region approximately 10 mm thick. 2D and 3D dynamic B_0_ shimming ROIs were chosen on the vessels within the 2D region (i.e., 1 slice with 2 mm slice thickness) and 3D region (i.e., 5 slices with 2 mm slice thickness each) centered at the labeling plane, respectively. Figures 4D and 4E show histograms depicting the frequency offset distribution of voxels within the target vessels in the 2D and 3D regions, respectively, before and after correction using global frequency offset, 2D, and 3D dynamic B_0_ shimming methods. Note that the corrected frequency offset values were based on simulated data, not actual measurements. The histograms clearly demonstrate a significant frequency offset before correction. While all methods successfully reduced off-resonance, 3D dynamic B_0_ shimming exhibited a narrower distribution of frequency offsets centered around 0 Hz (the target frequency offset) in the histogram, indicating improved field homogeneity and more effective shimming.

**Figure 4.**
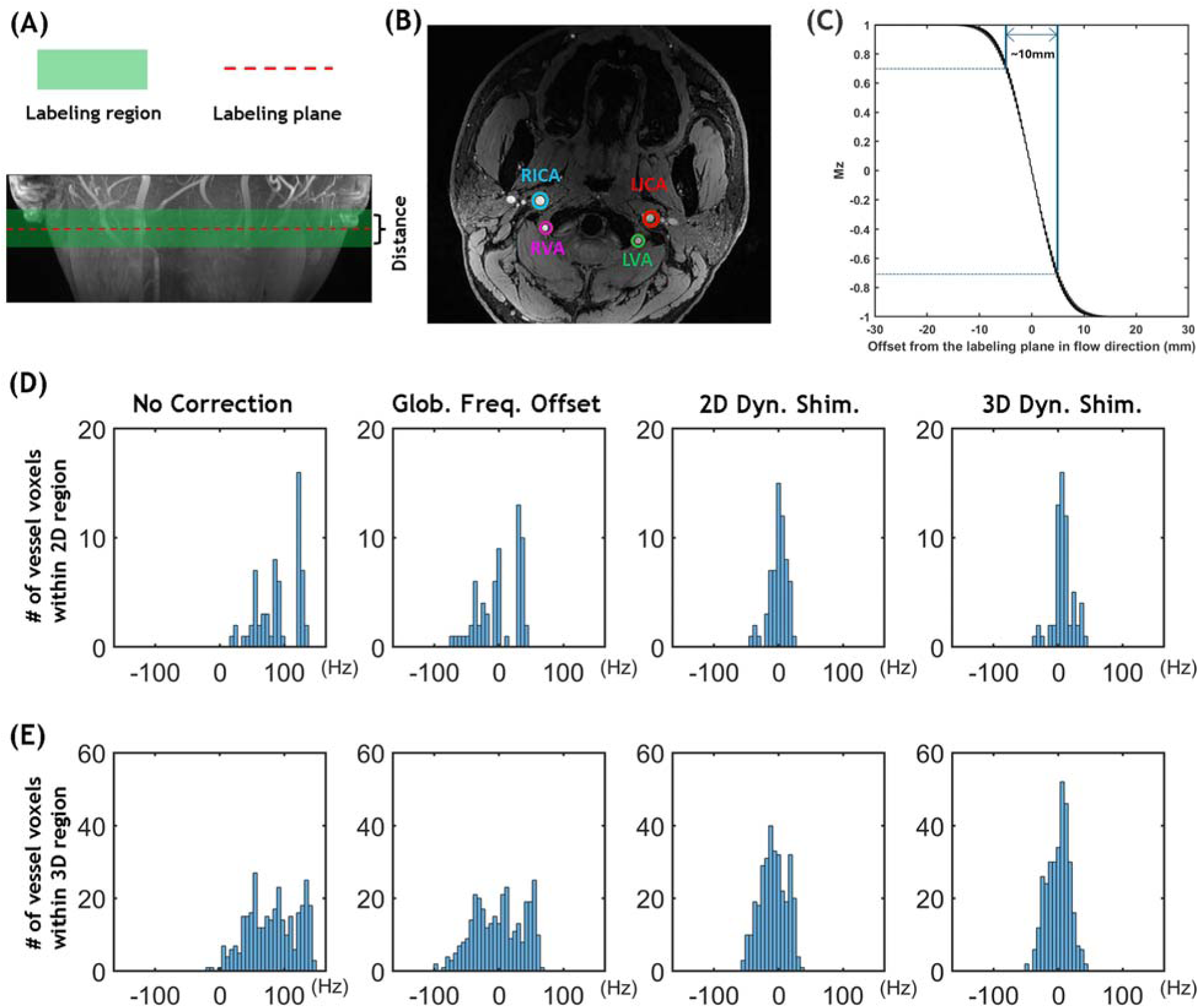
**(A)** The labeling region (green), where a flow-driven pseudo-adiabatic inversion process takes place in PCASL, and the labeling plane (black dashed line), also referred to as the central plane of the labeling region, are shown on a coronal TOF MIP. **(B)** TOF image at the labeling plane in the neck, highlighting the four main brain-feeding arteries (RICA, LICA, RVA, and LVA) with distinct colored circles. **(C)** The simulated evolution of the longitudinal magnetization (Mz) value during the PCASL labeling process. A threshold of ±70% was set to define the boundaries of the labeling region, resulting in a labeled area approximately 10 mm thick. Note that T_1_ and T_2_ relaxation times were set to be infinite in this simulation of PCASL to simplify interpretation. **(D, E)** Histograms of vessel voxel frequency offset distribution before and after global frequency offset correction, 2D and 3D dynamic B_0_ shimming, from a representative subject. **(D)** Distribution in 2D region (single slice at labeling plane). **(E)** Distribution in 3D region (five slices centered at labeling plane, representing the labeling region), demonstrating optimal field homogeneity with 3D dynamic B_0_ shimming.

Figure 5 compares the perfusion-weighted images from a volunteer obtained using PCASL without and with different off-resonance correction methods. The images were pair-wise subtracted and averaged following rigid registration, with no further masking or postprocessing applied. In the uncorrected acquisition, significant perfusion signal loss (indicated by red arrows) is evident in the posterior cerebral circulation due to off-resonance of the feeding arteries in the labeling region. This signal loss is substantially recovered in all the scans with application of the global frequency offset, OES-based, and 2D and 3D dynamic B_0_ shimming methods for correcting the off resonance. However, certain brain regions still exhibit lower signal intensity with the global frequency offset method compared to other correction methods (highlighted by yellow arrows). Notably, OES-based and 2D dynamic B_0_ shimming yielded similar signal intensity, while 3D dynamic B_0_ shimming exhibited the highest across all regions in this subject.

**Figure 5.**
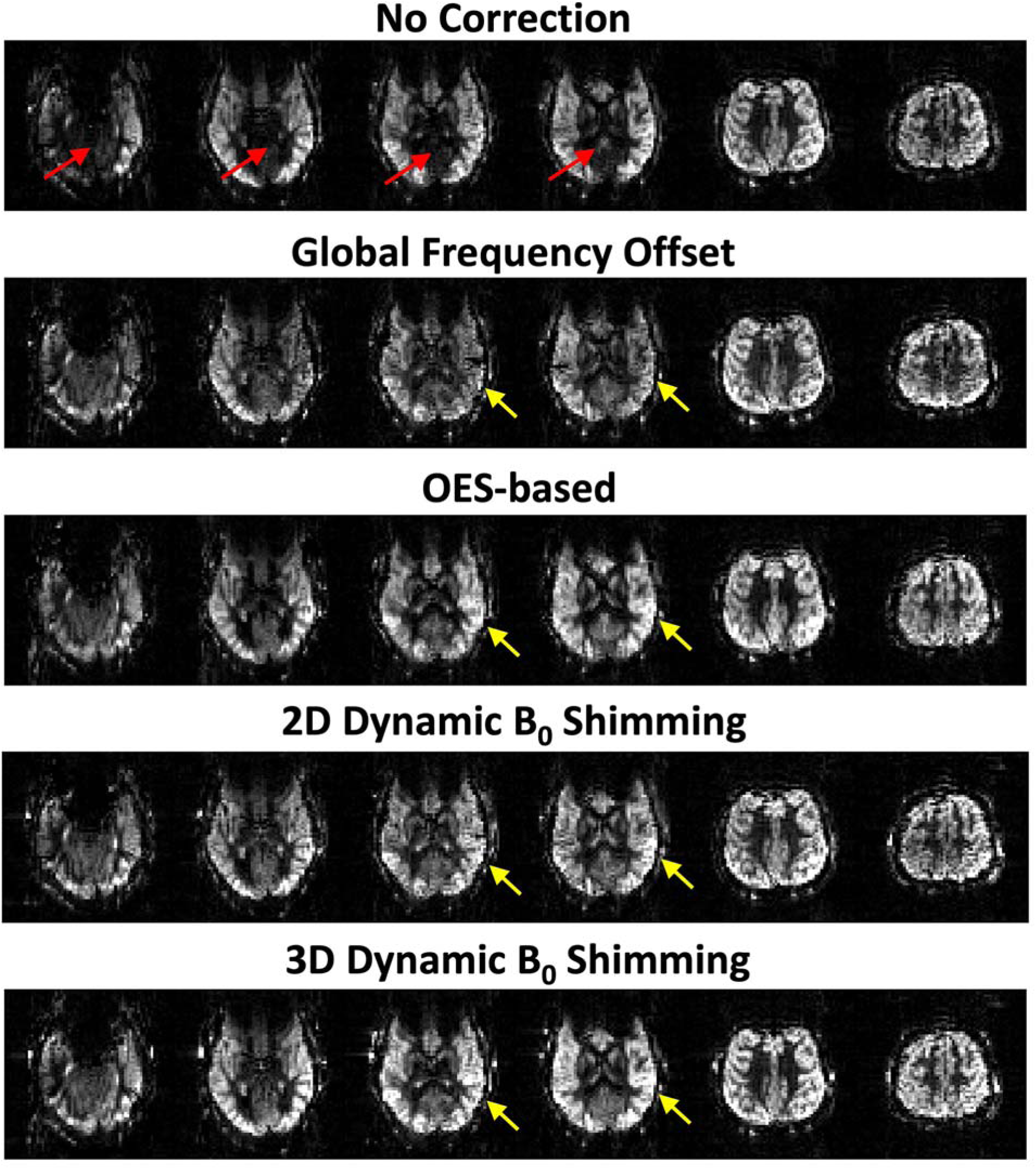
Comparison of whole-brain perfusion-weighted images acquired using PCASL with no correction, global frequency offset, OES-based, and 2D and 3D dynamic B_0_ shimming correction methods. In the uncorrected acquisition, significant perfusion signal loss (indicated by red arrows) is evident in the posterior cerebral circulation. The perfusion signal is improved after all correction methods, with differences in performance indicated by the yellow arrows, giving high quality perfusion images despite the short scan time (3.5 mins).

To emphasize the differences between 2D and 3D dynamic B_0_ shimming methods. Figure 6 presents a comparison of perfusion-weighted images using PCASL with both methods on the same volunteer (shown in Figure 5). Both methods produced high-quality whole-brain perfusion-weighted images. However, the 3D shimming method yielded an improved perfusion signal in the posterior circulation, as indicated by the red arrows. It can be inferred that the vertebral arteries exhibit better field homogeneity after 3D dynamic shimming compared to the 2D method.

**Figure 6.**
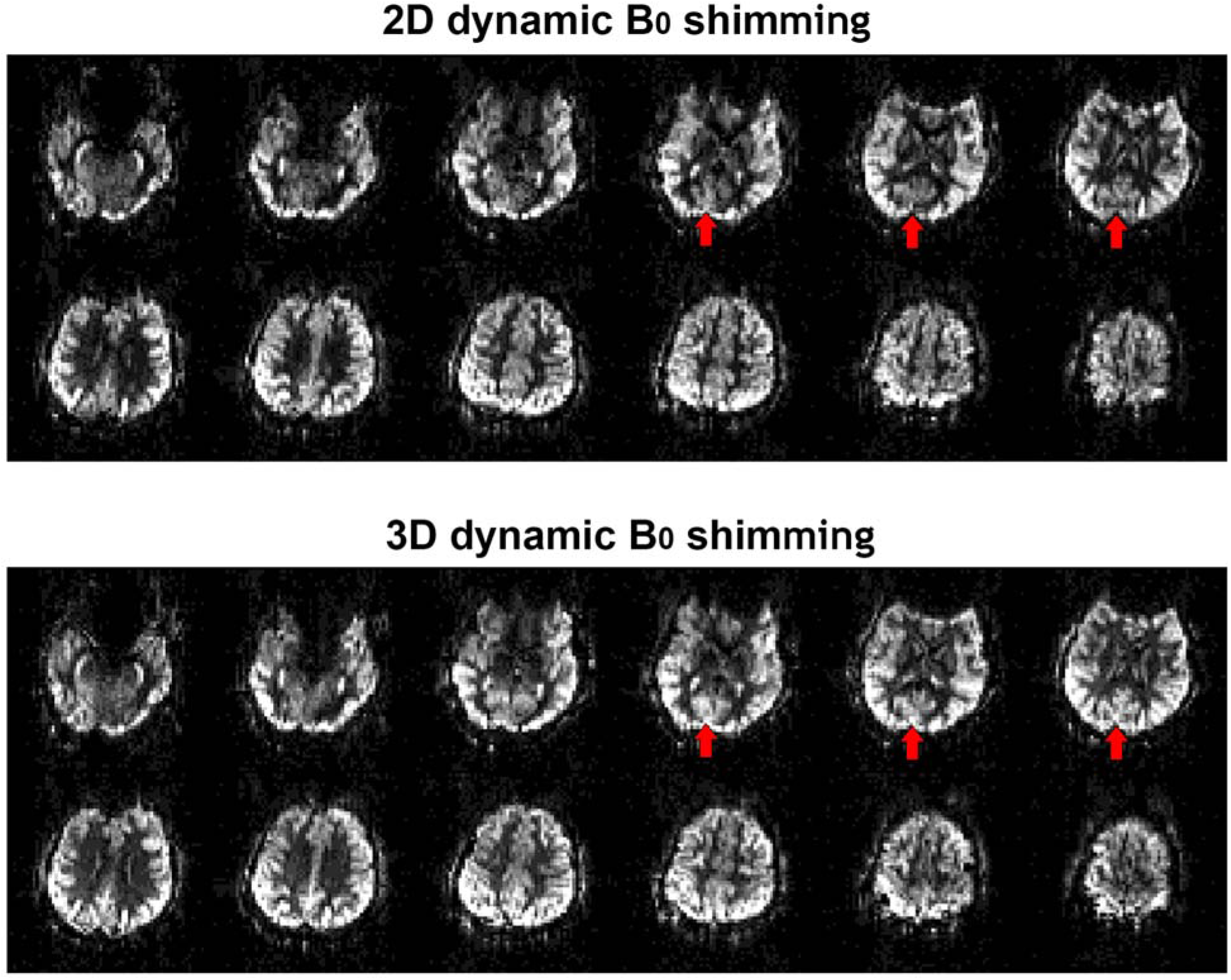
Comparison of whole-brain perfusion-weighted images acquired using PCASL with 2D dynamic B_0_ shimming and 3D dynamic B_0_ shimming methods on the same volunteer (shown in Figure 5). Enhanced perfusion signal can be observed in the posterior circulation when using the 3D dynamic B_0_ shimming method, as indicated by the red arrows.

Figure 7 compares the quantitative CBF values within the whole brain, white matter, and grey matter regions across the 6 volunteers. The results indicate that each method produces significantly improved perfusion values for the different brain regions. For the whole brain, the 3D dynamic B_0_ shimming method yields the highest perfusion value of 33.9±3 ml/100g/min, followed by the 2D dynamic B_0_ shimming and global offset methods, providing 31.5±3.2 and 29.9±2.6 ml/100g/min, respectively. The OES-based method provides the lowest value of 29.8±3.5 ml/100g/min. In grey matter, the 3D and 2D dynamic B_0_ shimming methods show the highest and second highest values of 42.9±6.7 and 40.5±7.9 ml/100g/min, respectively, while the global offset and OES-based methods yield similar but lower values of 38.2±7.1 and 38.2±6.5 ml/100g/min, respectively. In white matter, all four methods show a consistent pattern of 23.7±6.4 ml/100g/min, 20.3±5.4 ml/100g/min, 19.8±5.3 ml/100g/min, and 18.4±6.8 ml/100g/min. This consistent pattern suggests that the 3D dynamic B_0_ shimming method provides the best off-resonance correction and highest labeling efficiency, which results in higher perfusion values compared to the 2D dynamic B_0_ shimming, global offset, and OES-based methods.

**Figure 7.**
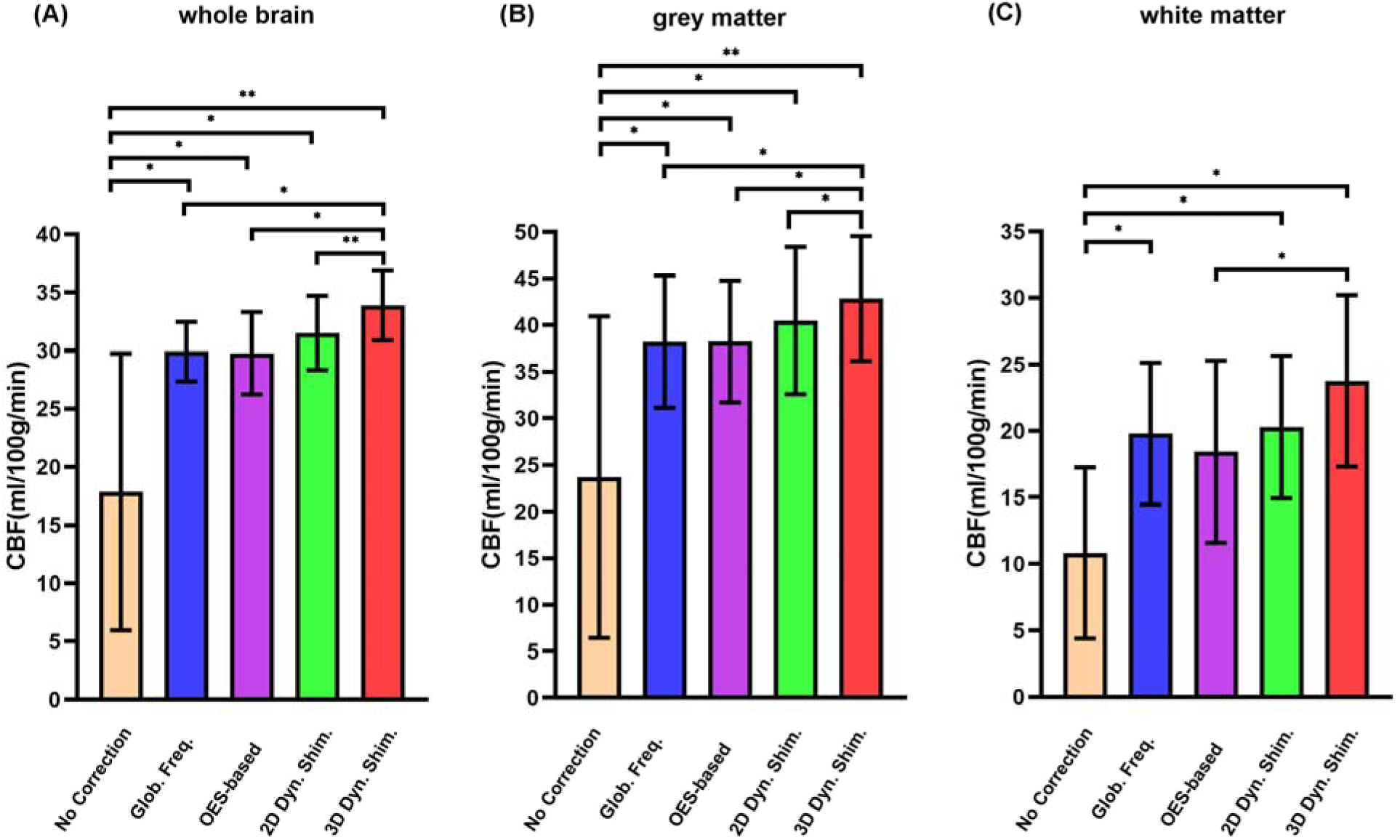
Quantitative CBF comparison across different correction methods. Average perfusion signal in **(A)** whole brain, **(B)** grey matter, and **(C)** white matter for six healthy subjects. Measurements were taken before off-resonance correction and after correction using the global frequency offset, OES-based, 2D dynamic B_0_ shimming, and 3D dynamic B_0_ shimming methods. (p-values are indicated by stars: * p<0.05, ** p<0.001).

To demonstrate the benefits of the improved SNR from UHF-PCASL, higher spatial resolution images were also acquired. Figure 8 demonstrates the ability of PCASL with 3D dynamic B_0_ shimming method at 7T to acquire high-resolution perfusion maps. Data from a representative subject is shown. The high-resolution maps provide finer anatomic details with a voxel size of 2.0 x 2.0 x 4 mm³, but at the cost of lower SNR compared to the low-resolution reference maps with a voxel size of 3.4 x 3.4 x 5 mm³. This comparison highlights the potential of PCASL at 7T for more detailed investigation of cerebral blood flow patterns in less than 5 minutes of scan time.

**Figure 8.**
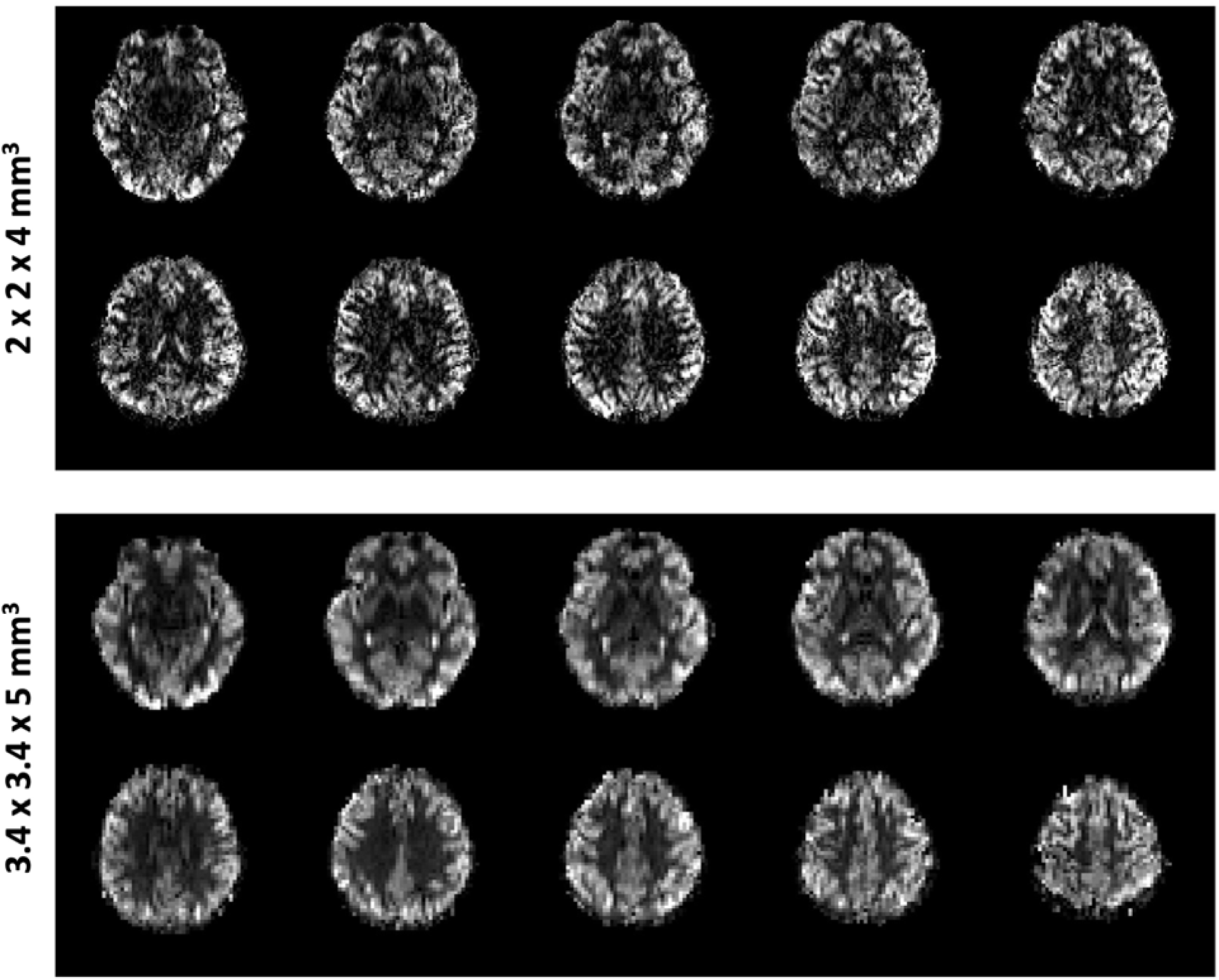
Demonstration of the feasibility of acquiring high spatial resolution (2.0 × 2.0 × 4 mm³) perfusion maps using PCASL with 3D dynamic B_0_ shimming at 7T, compared to low spatial resolution (3.4 × 3.4 × 5 mm³) maps from a representative subject. This comparison highlights the ability of PCASL at 7T to capture finer details of cerebral blood flow.

## Discussion

PCASL is recognized as the most preferred non-invasive perfusion technique among ASL variants at 3T (3,36–38). It employs a series of repeated low tip-angle RF pulses for tagging, which are feasible to implement on modern clinical MRI scanners, unlike the continuous RF pulses required for CASL, whilst also significantly mitigating magnetization transfer (MT) effects(7,10,39). Compared to 3T, PCASL at 7T should offer a very significant SNR increase, but in practice often exhibits a significant loss of perfusion signal due to off-resonance issues within the labeling region positioned far from the isocenter. To address this issue, our study introduces a novel dynamic B_0_ shimming method aimed at enhancing B_0_ field homogeneity, thereby improving the efficiency of arterial blood labeling.

Dynamic B_0_ shimming is also known as temporal control of the B_0_ shim fields or real-time B_0_ shimming, which was initially developed to improve 2D multi-slice MRI by customizing optimal shim settings for individual slices instead of settling for a global compromise (40,41). This method was further used for addressing time-varying B_0_ fluctuations caused by physiological processes like respiration(42–44), where the shim fields need to be updated in sync with the respiratory cycle. However, the proposed dynamic B_0_ shimming in this study only requires switching the shim settings between the labeling period and the rest of the sequence in PCASL. Since the standard gradient coils handle both imaging and shimming, linear dynamic B_0_ shimming can be achieved by manipulating the gradient amplitude directly, which has the same effect as adjusting the shim settings. Additionally, the gradients applied for shimming can only address the linear term of the field inhomogeneity by adding an offset to the linear gradients. Beyond this, there remains a residual global frequency offset and a nonlinear term of inhomogeneity. To compensate for the global frequency offset, we added an additional phase term to the PCASL RF pulses, addressing the additional off-resonance induced phase buildup between successive pulses. Whilst we found this to be effective in the experiments performed for this study, this approach does not account for the actual labeling plane being shifted by -2πΔf/(γG_max_). For example, with a 200 Hz offset (Δf) and a maximum gradient (G_max_) of 5.5 mT/m in the low SAR protocol, the labeling plane would shift by approximately 5 mm. An alternative method to compensate for the global frequency offset involves adjusting the central frequency (also known as the zero-order shimming parameter) during the labeling period by directly adding the Δf value to the central frequency. This would eliminate the need to modify the RF pulse phase and avoid the undesired labeling plane shift.

In this study, the labeling plane was chosen in the neck, where the four main brain-feeding arteries are located. Within this region, the arteries are relatively straight, and their geometry is simple. In such cases, the field inhomogeneity within the vessels can be well corrected by the linear terms using our targeted approach, with negligible contribution from nonlinear terms. However, PCASL and vessel-encoded PCASL(45) sometimes require placing the labeling plane above the Circle of Willis, a region rich in intricate vascular structures(46). Unfortunately, this proximity to the air-filled sinuses in the anterior of the brain significantly worsens field inhomogeneity. This, coupled with the need to label more vessels in this area, exacerbates higher-order field inhomogeneity, posing a substantial challenge for B_0_ shimming using only linear terms. Consequently, the proposed dynamic B_0_ shimming using the linear gradient would be unlikely to effectively address field inhomogeneity with significant nonlinear components. To handle this situation, higher-order spherical harmonic shimming coils may be utilized(47). However, dynamic shimming with these coils requires careful monitoring and compensation for the eddy currents induced by shim coil switching, a process not typically performed for second-order or higher-order shim coils on modern clinical scanners(48). Meanwhile, RF shimming coil arrays, which shape the B_0_ field by independently adjusting the direct current (DC), can also be employed for dynamic B_0_ shimming in PCASL (49,50). These arrays can be used separately or in combination with regular shimming coils.

It is valuable to compare our proposed dynamic B_0_ shimming method with others. Statistical analysis of the perfusion data in this study revealed that our 3D shimming method yielded significantly higher perfusion signal than all other evaluated methods. The 2D shimming method produces higher CBF than the OES-based and global frequency correction methods, although this difference is not statistically significant. In terms of sequence implementation, our proposed method preserves the sequence structure, making it simpler to integrate. The principles of 2D dynamic B_0_ shimming are theoretically similar to existing 2D methods like OES-based(16) method and OptPCASL(22), which utilize gradients blips between PCASL RF pulses to correct the linear component of in plane field inhomogeneity. However, our dynamic B_0_ shimming also includes the capability to apply gradients along the z-axis, addressing through-plane B_0_ variations, a capability not achievable with most other methods. Instead of using a simple shimming method that extends the static shimming region to the edge of the labeling region, a different approach can be proposed to shim the labeling and imaging regions separately. This method represents another type of dynamic B_0_ shimming. However, it shims the entire labeling region rather than just the vessel voxels within it, imposing greater constraints on the optimization process.

It is also worth mentioning that B_1_^+^ inhomogeneity is another major issue for 7T PCASL, which significantly affects the labeling efficiency. B_1_^+^ varies with head size and positioning, with a rapid drop in B_1_^+^ into the neck when using a head transmit coil at 7T and different B_1_^+^ distributions at the feeding arteries for each subject (51). In our study, we mitigate the B_1_^+^ drop by amplifying the RF flip angle by a certain factor. However, this approach did not address the fundamental issue of B_1_^+^ inhomogeneity across the labeled feeding arteries. In future studies, B_1_^+^ amplitude and its field homogeneity in the labeling plane can be improved by utilizing RF shimming with parallel transmission approaches or a dedicated labeling RF coil to enhance the labeling efficiency at 7T (52–55). It’s worth noting that the gradient-echo EPI readout used in this study may not be well-optimized for 7T imaging, as distortions and signal drop-out can be observed in the EPI images. To improve image quality, dynamic B_0_ shimming with optimal shimming parameters for each slice can be employed during the readout period. Additionally, non-EPI readout methods such as FLASH(19) can also enhance image quality. However, the improvements in perfusion signal achieved in this study should translate effectively to any readout method.

The current implementation of PCASL with dynamic B_0_ shimming involves several manual steps that hinder its clinical adoption. Crucial tasks, such as matching the static B_0_ shimming region between the field mapping scan and the PCASL scan, are performed manually, increasing the risk of human error. Additionally, the ROIs for the main feeding arteries within the dynamic B_0_ shimming region are chosen manually, and the parameters for dynamic B_0_ shimming are calculated offline. These manual processes complicate the PCASL setup and prolong scan time, making the technique less practical for routine clinical use. To address these limitations, the field mapping sequence for measuring off-resonance can be directly integrated into the PCASL scan as a pre-scan step. Furthermore, an automated ROI selection and dynamic B_0_ shimming calculation process should be developed and deployed directly on the scanner console. Integrating these features will significantly simplify the workflow for PCASL scans with dynamic B_0_ shimming, making the technique more efficient and user-friendly, promoting its wider application in clinical practice.

## Conclusion

Our study presents a novel dynamic B_0_ shimming approach that offers substantial benefits for enhancing PCASL at 7T. This method not only corrects in-plane linear B_0_ variations but also addresses the through-plane B_0_ variation within the labeling region. Importantly, our approach specifically targets B_0_ homogeneity within the vessels of interest within the labeling region, rather than the entire labeling area, allowing for more precise optimization compared to a generalized dynamic shimming approach. Overall, this rapid and comprehensive B_0_ shimming technique significantly improves the robustness and effectiveness of PCASL, unlocking the full potential of 7T ASL’s high sensitivity and spatial resolution for enhanced perfusion imaging.

## Code and Data Availability Statement

The code for the dynamic shimming calculations can be found at <LINK to be added upon publication>.

We are currently unable to share subject level data due to data protection issues, although our centre is actively working on a solution to this. Data underlying the plots in this paper can be found at <LINK to be added upon publication>.

## Acknowledgments

This study was supported by the Oxford NIHR Biomedical Research Centre. The Wellcome Centre for Integrative Neuroimaging is supported by core funding from the Wellcome Trust (203139/Z/16/Z). T.W.O. is supported by a Sir Henry Dale Fellowship jointly funded by the Wellcome Trust and the Royal Society (220204/Z/20/Z). We are grateful to Yulin Chang for providing example code showing how to change linear shim terms on a Siemens platform. For the purpose of open access, the author has applied a CC BY public copyright license to any Author Accepted Manuscript version arising from this submission.

## Notes

### Competing Interest Statement

The authors have declared no competing interest.

